# Loss of LGR5 through plasticity or gene ablation is associated with therapy resistance and enhanced MET-STAT3 signaling in colorectal cancer cells

**DOI:** 10.1101/2022.03.01.482539

**Authors:** Tressie A. Posey, Joan Jacob, Ashlyn N. Parkhurst, Shraddha Subramanian, Liezl E. Francisco, Zhengdong Liang, Kendra S. Carmon

**Author notes:** **Corresponding Author:** Kendra S. Carmon, Center for Translational Cancer Research, The Brown Foundation Institute of Molecular Medicine, University of Texas Health Science Center at Houston, 1825 Pressler St., Houston, TX 77030.

## Abstract

Plasticity plays a significant role in colorectal tumor initiation, progression, and drug resistance. LGR5 is highly expressed in colorectal cancer (CRC) and marks functional cancer stem cells (CSCs). While LGR5^+^ CSCs are tumor-initiating, the majority of CRC cells that disseminate to seed metastases are LGR5^-^; however, reemergence of LGR5^+^ CSCs is required to drive metastatic outgrowth. LGR5^+^ CSCs have been shown to convert to LGR5^-^ CRC cells in response to chemotherapies and this loss of LGR5 promotes a more drug-resistant phenotype. However, the molecular mechanisms that mediate plasticity remain elusive. In this study, we demonstrate conversion of LGR5^+^ CRC cells to an LGR5^-^ state in response to chemotherapy, LGR5-targeted antibody-drug conjugates (ADCs), or LGR5 gene ablation, led to activation of STAT3. Further investigation revealed increased STAT3 activation occurred a result of increased MET activity. LGR5 overexpression decreased MET-STAT3 activity and sensitized CRC cells to therapy. STAT3 inhibition suppressed MET phosphorylation, while constitutively active STAT3 reduced LGR5 levels and increased MET activity, suggesting a potential feedback mechanism. Combination treatment of MET-STAT3 inhibitors with irinotecan or ADCs substantiated synergistic effects in vitro. In CRC xenografts, STAT3 inhibition combined with irinotecan enhanced tumor growth suppression and prolonged survival. These findings suggest a mechanism by which drug-resistant LGR5^-^ CRC cells acquire a survival advantage through activation of MET-STAT3 and provide rationale for new treatment strategies that target CRC cell plasticity.

**Significance:** This study reveals that transition of highly plastic LGR5^+^ CRC cells to a more drug-resistant LGR5^-^ state involves activation of MET-STAT3 signaling and provides new insight into therapeutic strategies to combat plasticity.

## Introduction

Colorectal cancer stem cells (CSCs), or tumor-initiating cells, mediate drug resistance and relapse through the capacity to either self-renew or differentiate into heterogeneous lineages of tumor cells (1,2). LGR5, or Leucine-rich repeat containing, G protein-coupled Receptor 5 (LGR5), is highly expressed in approximately 60-70% of CRCs (3-5) and is a validated marker of normal adult intestinal stem cells and functional CSCs (6-8). Originally identified as a Wnt target gene, LGR5 and its related receptors LGR4 and LGR6 bind the R-spondin (RSPO1-4) growth factors responsible for modulating Wnt/β-catenin signaling, an important pathway implicated in CRC and stem cell maintenance (9-12).

LGR5^+^ CSCs can initiate colorectal tumor growth and exhibit a high level of plasticity, or the capacity to shift between CSC and non-CSC states during tumor progression and to circumvent therapy. Elimination of LGR5^+^ CSCs in CRC tumors using either genetic ablation approaches or LGR5-targeted antibody-drug conjugates (ADCs) resulted in tumor inhibition or stasis, with relapse after treatment termination (3,5,6,8). LGR5^-^ CRC cells were able to sustain tumors, however the reemergence of LGR5^+^ CSCs was proven necessary for tumor regrowth. More recently, it was reported that the majority of CRC cells that disseminate from the primary tumor for seeding liver metastases are LGR5^-^ (13). However, conversion of LGR5^-^ CRC cells to an LGR5^+^ state was necessary to promote metastatic outgrowth, demonstrating the dynamic plasticity that exists during disease progression and metastasis. Furthermore, treatment with standard chemotherapies was shown to trigger LGR5^+^ CSCs to transition to an LGR5^-^, drug-resistant state (14). Correspondingly, we reported that knockdown or knockout of LGR5 in CRC cell lines also conferred a more drug-resistant phenotype (15). Though LGR5^+^ CSC plasticity is well established, the molecular and cellular processes underpinning plasticity remain relatively unknown.

In this study, we aimed to identify a molecular mechanism involved in mediating plasticity and drug resistance in LGR5^-^ CRC cells. We show that conversion of CRC cells from LGR5^+^ to LGR5^-^ state in response to different therapies or LGR5 gene ablation leads to activation of the MET-STAT3 pathway. Our results further suggest that LGR5^-^ CRC cells acquire a survival advantage through MET-STAT3 activation to become more resistant to treatment. Combination therapy of MET-STAT3 pathway inhibitors with standard chemotherapy enhanced treatment efficacy and prolonged survival. These findings identify LGR5 as a novel regulator of MET-STAT3 signaling and may lead to new treatment strategies for targeting plasticity of CRC cells.

## Methods and Materials

### Chemicals and plasmids

The anti-LGR5 ADC (anti–LGR5-mc-vc-PAB-MMAE) with drug-to-antibody ratio of 4 was generated as previously described (3). Irinotecan and 5-fluorouracil (5-FU) were purchased from Selleck and Acros Organics, respectively. Stattic was purchased from Tocris, Cryptotanshinone and Gefitinib were from Selleck, Crizotinib from Cell Signaling, and XAV939 from Cayman Chemical. The plasmid encoding myc-LGR5 was generated previously (9). Constitutively active mouse STAT3 (STAT3-CA) plasmid Stat3-C Flag pRc/CMV was a gift from Jim Darnell (Addgene plasmid 8722).

### Cell Culture

LS180, HCT116, and DLD-1 colorectal cancer cells were purchased from the ATCC. LoVo colorectal cancer cells were obtained from Dr. Shao-Cong Sun (M.D. Anderson Cancer Center). Cell lines were authenticated utilizing short tandem repeat profiling, routinely tested for mycoplasma, and cultured in RPMI medium supplemented with 10% FBS and penicillin/streptomycin at 37°C with 95% humidity and 5% CO_2_. Transient transfections were performed using jetPRIME (Polypus Transfection). Stable CRISPR/Cas9 LGR5 knockout clonal LS180 cell lines 1.4 and 1.5 were generated using the lenti-CRISPRv2 vector system as we described (15). Stable shRNA LGR5 knockdown LoVo cell lines pLKO.1 (shCTL), shLGR5-1, and shLGR5-2 were generated as we previously reported (3,10).

### Western blot

For western blot analysis, protein extraction was performed using RIPA buffer (Sigma) supplemented with protease/phosphatase inhibitors. Cell lysates were incubated at 37°C for 1 hour in Laemmli SDS sample buffer prior to loading on SDS-PAGE. Commercial antibodies used in this study were: anti-LGR5 (ab75732), anti-EGFR Y1173 (ab32578) from Abcam and anti-p-STAT3 Y705 (9145), anti-STAT3 (9139), anti-p-MET Y1234/1235 (3077), anti-MET (8198), anti-p-EGFR Y1068 (3777), anti-EGFR (54359), anti-non-p-β-catenin (8814), anti-β-catenin (8480), anti-Cyclin D1 (55506), anti-Bcl-xl (2764), anti-flag (14793), and anti-β-actin (3700) from Cell Signaling. Horseradish peroxidase-labeled anti-mouse (Cell Signaling, 7076) and anti-rabbit (Cell Signaling, 7074) secondary antibodies were utilized for detection with the standard ECL protocol.

### Cell viability assays

Cell viability assays were performed as previously described (15) using CellTiter-Glo® luminescent cell viability assay (Promega). Serial dilutions of commercial drugs or anti-LGR5 ADC were added at the indicated concentrations and allowed to incubate at 37°C for 3 to 4 days. IC50s were determined using Prism 5 (GraphPad Software, Inc.). Representative results of at least 3 independent experiments are shown.

### Clonogenicity assays

For colony formation assays, cells were seeded in 6-or 12-well plates at 5000 or 1000 cells per well, respectively. Experiments were performed at least three times and colonies were quantified by manual counting or using ImageJ. For soft agarose assays, cells were seeded in culture medium containing low melting point agar at a density of 500 cells per well in 12-well plates. After 2 weeks, colonies were stained with crystal violet and the numbers of colonies were counted manually or quantified using ImageJ.

### Immunocytochemistry

Cells were seeded into 8-well chamber slides and allowed to adhere overnight. For LGR5 internalization experiments, cells were treated with irinotecan at the indicated time points and incubated with anti-LGR5 rh8F2 mAb (16) at 37°C for 30 minutes. Cells were washed, fixed in 4% formaldehyde, permeabilized with 0.1% saponin (Sigma), and incubated with anti-human-Alexa 555 (Invitrogen) for 1 hour at room temperature. For detection of STAT3 translocation, cells were fixed, permeabilized with 0.3% 1X Triton-X and incubated with anti-STAT3 Ab (Cell Signaling, 9139), followed by anti-mouse-Alexa-488 (Invitrogen). Nuclei were counterstained with TO-PRO®-3 (Invitrogen). Images were acquired using confocal microscopy (Leica TCS SP5 microscope) with the LAS AF Lite software (Leica Microsystems, Inc.).

### Animal Studies

In vivo experiments were carried out in strict accordance with the recommendations of the Institutional Animal Care and Use Committee of the University of Texas Health Science Center at Houston (AWC-20-0144). Female 6-to 8-week-old nu/nu mice (Charles River Laboratories or Jackson Laboratory) were subcutaneously inoculated with 1 × 10^6^ LoVo or 2 × 10^6^ LS180 cells in 1:1 mixture of PBS and matrigel (BD Biosciences) into lower right flank. Once tumors reached an average size of ∼100-150 mm^3^, animals were randomized into four treatment groups (vehicle, stattic, irinotecan, or irinotecan + stattic). Irinotecan treatment was initiated on Day 0 and stattic treatment was initiated on Day 2. Irinotecan (20 mg/kg) or vehicle of 2% DMSO in 5% dextrose in sterile water. was administered intraperitoneally every five days (LoVo = 3 doses; LS180 = 2 doses). Stattic (10 mg/kg) or vehicle was administered intraperitoneally every other day (LoVo = 6 doses; LS180 = 3). Tumor volumes were measured at least biweekly and estimated by the formula: Tumor volume = (length × width^2^)/2. Mice were euthanized when tumor volume reached approximately 2000 mm^3^. Once the first tumor from each vehicle group reached maximum tumor burden, treatment was terminated, and animals were monitored for changes in survival.

### Statistical Analysis

Statistical analysis was performed and IC50s determined using the Prism 5 (GraphPad Software, Inc.). All in vitro experiments were performed at least three times. The levels of significance between samples were determined through an unpaired two-tailed Student t test (mean comparison with one factor) or one-way ANOVA for groups with multiple comparisons. Combination index (CI) values were calculated by the Chou-Talalay method using CompuSyn (17). For in vivo experiments, statistical significance of differences on tumor growth was determined using one-way ANOVA and Dunnett’s multiple comparison test and the Log-rank (Mantel-Cox) test was used for survival studies. Data are shown as mean ± standard error of the mean (SEM) or standard deviation (SD) as indicated, with P values ≤ 0.05 considered to be statistically significant.

### Data and Materials Availability

All data are contained within the article and materials can be requested from the corresponding author upon reasonable request.

## Results

### Drug-induced plasticity of CRC cells promotes a resistant LGR5-negative state

We previously showed that loss of LGR5 expression increased drug resistance to chemotherapies and proliferation of CRC cells (15). Furthermore, treatment with LGR5-targeted ADCs eliminated LGR5^+^ CRC tumors, however a fraction of tumors relapsed with LGR5-low/negative expression (3). Therefore, we questioned how both gene ablation and treatment-induced loss of LGR5 through plasticity promote a more drug-resistant state and if similar molecular mechanisms are involved. First, we tested the effect of chemotherapy and targeted ADCs on LGR5 expression to verify that LGR5^+^ CRC cells convert to an LGR5^-^ state during drug treatment. LoVo and LS180 CRC cell lines were selected based on their high endogenous expression of LGR5 (15). CRC cells were treated with 10 µM irinotecan or 6.5 nM (or 1 µg/ml) anti-LGR5 ADC (anti-LGR5-mc-vc-PAB-MMAE) at the indicated time-points and changes in LGR5 expression were measured by western blot (Figs. 1A-B). LGR5 was nearly undetectable after 72 hours of treatment in both LoVo and LS180 cells. Comparable results were observed after treatment with 10 µM 5-fluoruoracil (Supplementary Fig. S1C). Immunocytochemistry (ICC) staining revealed that LGR5 is homogenously expressed throughout the CRC cell lines and verified that transition from an LGR5^+^ to an LGR5^-^ state is due to drug-induced plasticity rather than survival of cells that originated as LGR5^-^. Comparable to western analysis, irinotecan treatment led to an approximate 75% and 85% decrease in LGR5^+^ LoVo and LS180 cells, respectively, after 24 hours; with near complete loss in LGR5^+^ expression after 72 hours (Fig. 1C). As shown in Fig. 1D, LGR5^-^ CRC cells transition back to an LGR5+ state after irinotecan and ADC washout, providing further evidence of plasticity. These data demonstrate drug-induced CRC plasticity and suggest that the LGR5^-^ state may be more resistant due to activation of survival signaling mechanisms.

**Figure 1.**
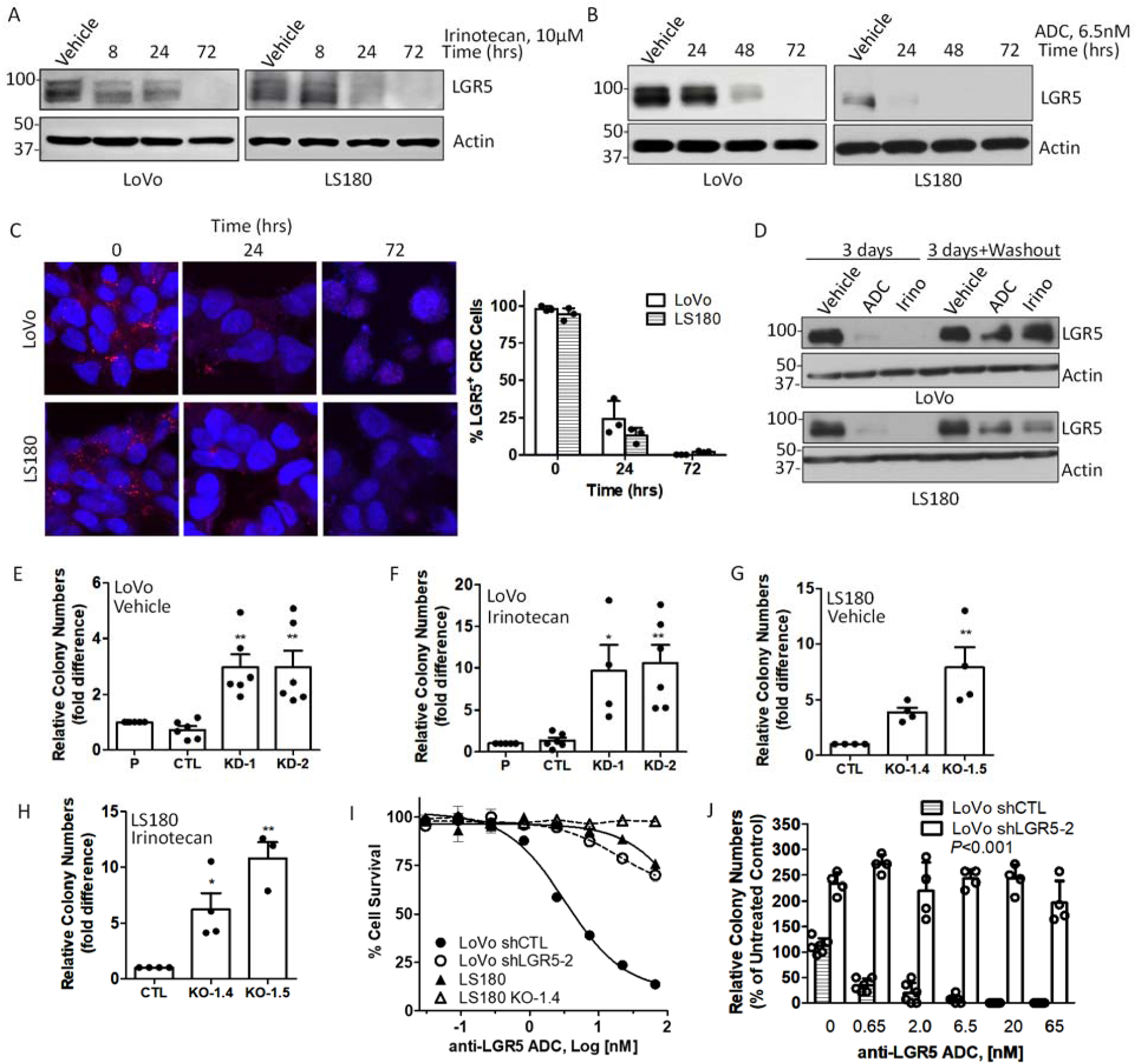
Treatment-induced loss or gene ablation of LGR5 converts CRC cells to a more drug-resistant state. (A-B) Western blot showing time course of changes in LGR5 expression in LoVo and LS180 CRC cells treated with (A) irinotecan (10 µM) or (B) anti-LGR5 ADC (6.5 nM). (C) Confocal microscopy images of LGR5 expression in LoVo and LS180 after inrinotecan (10 µM) treatment. (D) Western blot of LGR5 expression in LoVo and LS180 cells treated with anti-LGR5-ADC (6.5 nM) or irinotecan (10 µM) for 72 hours or treated with drugs followed by washout and allowed to recover for 72 hours. (E-H) Quantification of colony formation assays performed in LoVo shRNA control and LGR5 knockdown (KD) cells treated with (E) vehicle or (F) irinotecan (10 µM) and LS180 control and LGR5 knockout (KO) cells treated with (G) vehicle or (H) irinotecan (5 µM). (I) Cytotoxicity of anti-LGR5 ADC on CRC cell lines after 4 days. (J) Quantification of anti-LGR5 ADC dose-dependent effects on LoVo control and LGR5 KD cells using the soft agar colony formation assay. Experiments were performed at least 3 times. Statistical analysis was performed using one-way ANOVA. *, P ≤ 0.05, **, P ≤ 0.01, compared to controls. Error bars are SEM.

### Deletion of LGR5 enhances survival and drug resistance

As altered survival capacity of CRC cells is a critical determinant of drug resistance and metastatic potential of tumor cells, we examined changes in clonogenicity with loss of LGR5. We previously generated two LoVo LGR5 knockdown (KD) cell lines with independent shRNAs (shLGR5-1 and shLGR5-2) and two LS180 LGR5 knockout (KO) cell lines using the CRISPR/Cas9 system (1.4 and 1.5) (15). Clonogenicity assays were performed in the absence and presence of irinotecan for LoVo and LS180 cell lines (Figs. 1E-H). LGR5 KD/KO cells showed increased colony formation compared to parental (P) and vector control cell lines (CTL) when seeded at low density, approximately 3-fold for LoVo LGR5 KD cells and 6-to 11-fold for LS180 LGR5 KO cells (Figs. 1E,1G, and Supplementary Fig. S1B-C). The enhanced resistance and clonogenic capacity of LGR5 KD/KO cells was further demonstrated when cells were subjected to irinotecan. Clonogenicity of LoVo LGR5 KD (10-fold) and LS180 LGR5 KO cells (4-to 7-fold) was significantly increased compared to respective controls (Figs. 1F,1H, and Supplementary Figs. S1B-C). Similar findings were also shown in the presence of 5-fluoruoracil (Supplementary Figs. S1D). We reported that LoVo cells were sensitive to anti-LGR5 ADCs linked to an MMAE payload, however other CRC cell lines are more resistant to MMAE, rendering MMAE conjugated ADCs less effective despite high LGR5 expression. As shown in Fig. 1I, LoVo cells respond to ADC treatment with an approximate IC_50_ = 2-5 nM, whereas LS180 cells are relatively resistant. This may be attributed to a 10-fold difference in MMAE potency for LS180 compared to LoVo cells (IC_50_ = 2.3 vs. 0.2 nM, respectively; Supplementary Fig. S1E). ADC efficacy was diminished in both LGR5 KD and KO cell lines, demonstrating ADC specificity (Fig. 1I). Treatment of LoVo cells with ADC reduced clonogenicity in soft agar in a dose dependent manner (Fig. 1J and Supplementary Fig. S1F). As expected, LGR5 KD increased clonogenicity approximately 2.5-fold in soft agar and cells were resistant to LGR5-targeted ADC treatment (Fig. 1J and Supplementary Fig. S1F). These results demonstrate that loss of LGR5 expression increases survival and clonogenicity in vitro.

### LGR5 KD/KO in CRC cells increases STAT3 activation mediated through Met

To identify signaling mechanisms that may mediate plasticity and drug resistance in LGR5 KD/KO CRC cells, we performed a western blot analysis to determine changes in protein expression. In LoVo cells, LGR5 KD resulted in increased levels of active (non-phosphorylated) β-catenin, consistent with our previous report (10). Interestingly, we also detected increased phosphorylation of STAT3 with LGR5 KD (Fig. 2A). Increased nuclear accumulation of p-STAT3 was detected by ICC in LoVo LGR5 KD cells (Fig. 2B), suggesting increased STAT3-mediated transcriptional activity. Consistently, we also observed changes in the expression levels of STAT3-associated survival proteins involved in proliferation and inhibition of apoptosis (18). Specifically, loss of LGR5 resulted in increased Cyclin D1 and Bcl-xL expression (Fig. 21). These findings show that loss of LGR5 can increase both Wnt/β-catenin and STAT3 signaling in CRC cells.

**Figure 2.**
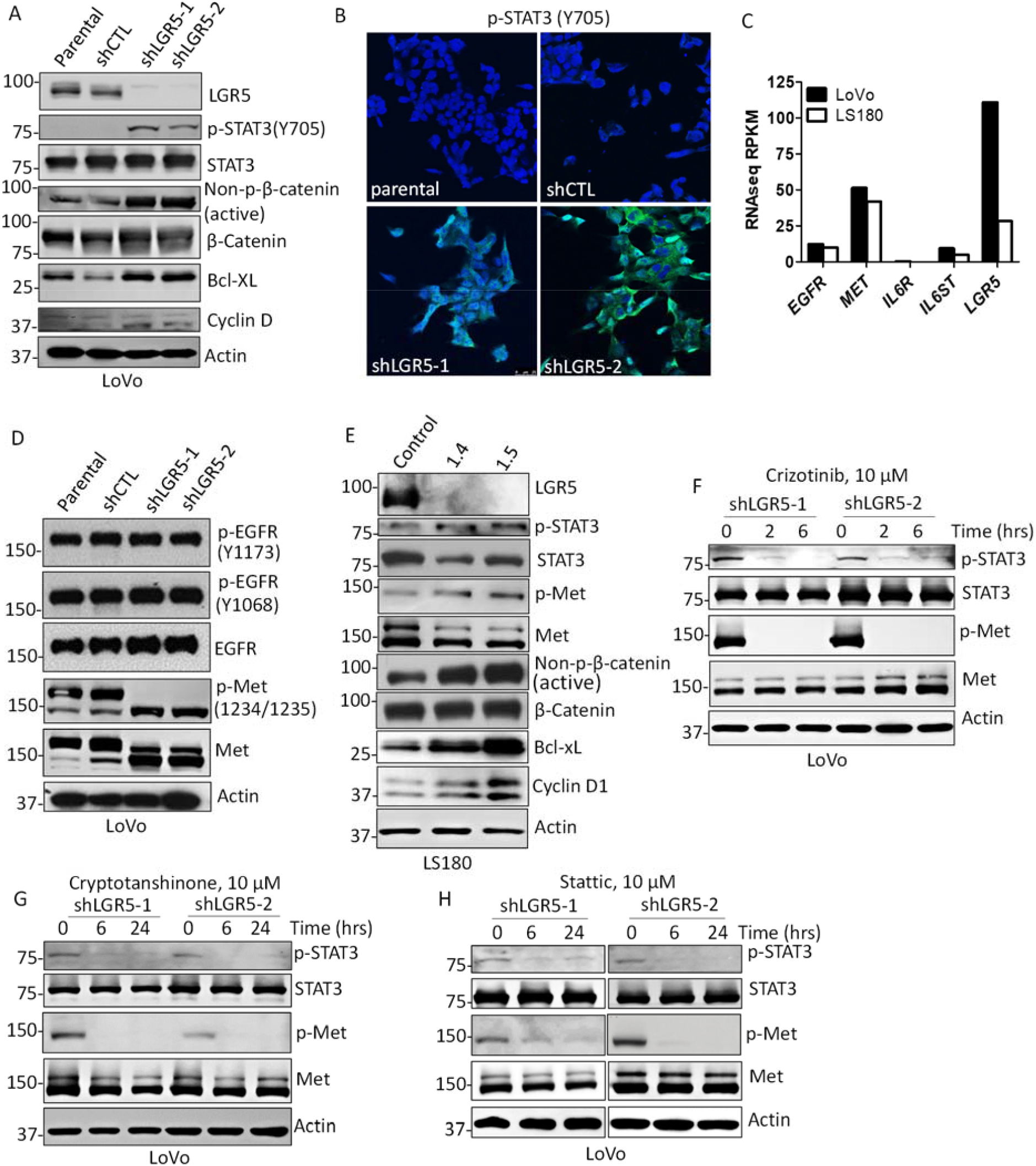
Loss of LGR5 enhances phosphorylation of STAT3 through activation of MET. (A) Western blot of changes in β-catenin, STAT3, and associated target proteins in LoVo control and LGR5 KD cells. (B) Confocal microscopy images of phosphorylated STAT3 in LoVo cell lines. (C) RNAseq expression data for LoVo and LS180 cells from CCLE. (D) Western blot of EGFR and MET (β-subunit) in LoVo cell lines. (E) Western blot of changes in β-catenin, STAT3, MET and associated target proteins in LS180 control and LGR5 KO cells. (F) Effect of vehicle (0.1% DMSO), MET inhibitor (crizotinib, 10 μM), or STAT3 inhibitor (stattic, 10 μM) on phosphorylation in LoVo LGR5 KD and LS180 LGR5 KO cells after 6 hours. Experiments were performed at least 3 times.

To identify which upstream mediators may be responsible for increased STAT3 phosphorylation, we characterized changes in expression and activation of membrane receptors known to mediate STAT3 activation, including IL6R/Gp130, EGFR, and MET (19-21). RNAseq data retrieved from the Cancer Cell Line Encyclopedia (CCLE) database showed that LoVo and LS180 CRC cells do not express *IL6R* and have minimal expression of *IL6ST* (Gp130) (Fig. 2C). We also failed to detect IL6R by western blot in LoVo cells (not shown). Compared to *LGR5* expression, both cell lines express high levels of *MET* and moderate expression levels of *EGFR* (Fig. 2C). To determine if Gp130 was playing a role in STAT3 activation, LGR5 KD LoVo cells were treated with Sc144, a small molecule Gp130 inhibitor (Supplementary Fig. S2A). However, inhibition of Gp130 failed to suppress STAT3 activation, suggesting that IL6R/Gp130 is not responsible for aberrant STAT3 signaling. We next examined LoVo LGR5 KD cell lines for potential changes in activation of EGFR and MET by western blot. As shown in Fig 2D, no significant changes in EGFR phosphorylation at sites known to mediate STAT3 signaling (Y1068 or Y1173) were observed with LGR5 KD. Furthermore, LoVo LGR5 KD cells treated with the EGFR inhibitor, gefitinib, did not result in decreased STAT3 activation (Supplementary Fig. S2B). These findings suggest that STAT3 activation in LGR5 KD CRC cells is most likely not mediated through IL6R/Gp130 or EGFR activation.

The MET receptor tyrosine kinase (RTK), initially synthesized as a partially glycosylated 170 kDa single-chain intracellular precursor (pro-Met), undergoes extensive posttranslational modifications to become a functionally mature protein (22). Mature MET is composed of a 45 kDa α-subunit that is linked by two disulfide bonds to a 145 kDa β-subunit (22). Of note, LoVo cells exhibit a defect in the normal processing of pro-MET, leading to a 190 kDa single-chain protein (23). Interestingly, in LGR5 KD cells we observed an increase in both the total and phosphorylated levels of the processed, mature form of MET as indicated by the 145-kDa band (Fig. 2D). Similarly, we detected an approximate 2-fold increase in phosphorylation of MET and STAT3 and increased levels of active β-catenin in LS180 LGR5 KO and DLD-1 LGR5 KD cells (Fig. 2E and Supplementary Fig. S2C). Similar to LoVo LGR5 KD cells, LS180 LGR5 KO cells had increased protein expression of STAT3-associated target genes Cyclin D1 and Bcl-xL (Fig. 2E). The effect of MET inhibition on STAT3 was investigated by treating LGR5 KD/KO cells with the MET inhibitor crizotinib, which functions through competitive binding within the ATP binding pocket (Fig. 2F and Supplementary Fig. S2D). Western blot showed that crizotinib treatment resulted in loss of both phosphorylated MET and STAT3 (Figs. 2F and Supplementary Fig. S2D), suggesting that STAT3 activation is mediated through MET. We also observed loss of active MET after treatment of LGR5 KD/KO cells with STAT3 inhibitors stattic and cryptotanshinone (Fig. 2F and Supplementary Figs. S2E-F), which inhibit STAT3 by interacting with the SH2 domain, preventing Tyr705 phosphorylation and dimerization (24,25). Of note, stattic treatment also resulted decreased levels of total STAT3 in LS180 cells. STAT3 inhibition of MET indicates the involvement of a potential feedback mechanism between MET and STAT3. These data suggest that elevated STAT3 signaling in LGR5 KD/KO CRC cells is mediated through increased MET activation.

### Inverse regulation of LGR5 and STAT3

To further elucidate the relationship between LGR5 expression and MET-STAT3 activation, constitutively activate STAT3 (STAT3-CA) was overexpressed in both LoVo and LS180 cells by transient transfection. Western blot showed that STAT3-CA concomitantly reduced LGR5 expression levels and increased MET phosphorylation (Fig. 3A). Cell viability assays performed using CellTiter-Glo showed that STAT3-CA cells were more resistant to irinotecan and anti-LGR5 ADCs, supporting a role for STAT3 in mediating resistance in LGR5 KD/KO CRC cells (Figs. 3B-D). Average IC_50_s values increased approximately 2 to 3-fold with STAT3-CA overexpression in both LoVo and LS180 cells. We next tested if overexpression of LGR5 could modulate MET-STAT3 activation. We selected the HCT116 CRC cell line, as it does not endogenously express LGR5 and has higher baseline levels of phosphorylated STAT3 compared to LoVo and LS180 cells (Supplementary Fig. S2G). LGR5 was overexpressed by transient transfection using increasing amounts of DNA (Fig 3D). Western blot showed gradual loss of MET and STAT3 activation with increasing amounts of transfected LGR5 (Fig. 3E). LGR5 overexpression also sensitized HCT116 cells to irinotecan treatment, reducing the IC_50_ by approximately 5-fold (Fig. 3F). These findings further support a role for MET-STAT3 signaling in driving drug resistance and survival in CRC cells with loss of LGR5.

**Figure 3.**
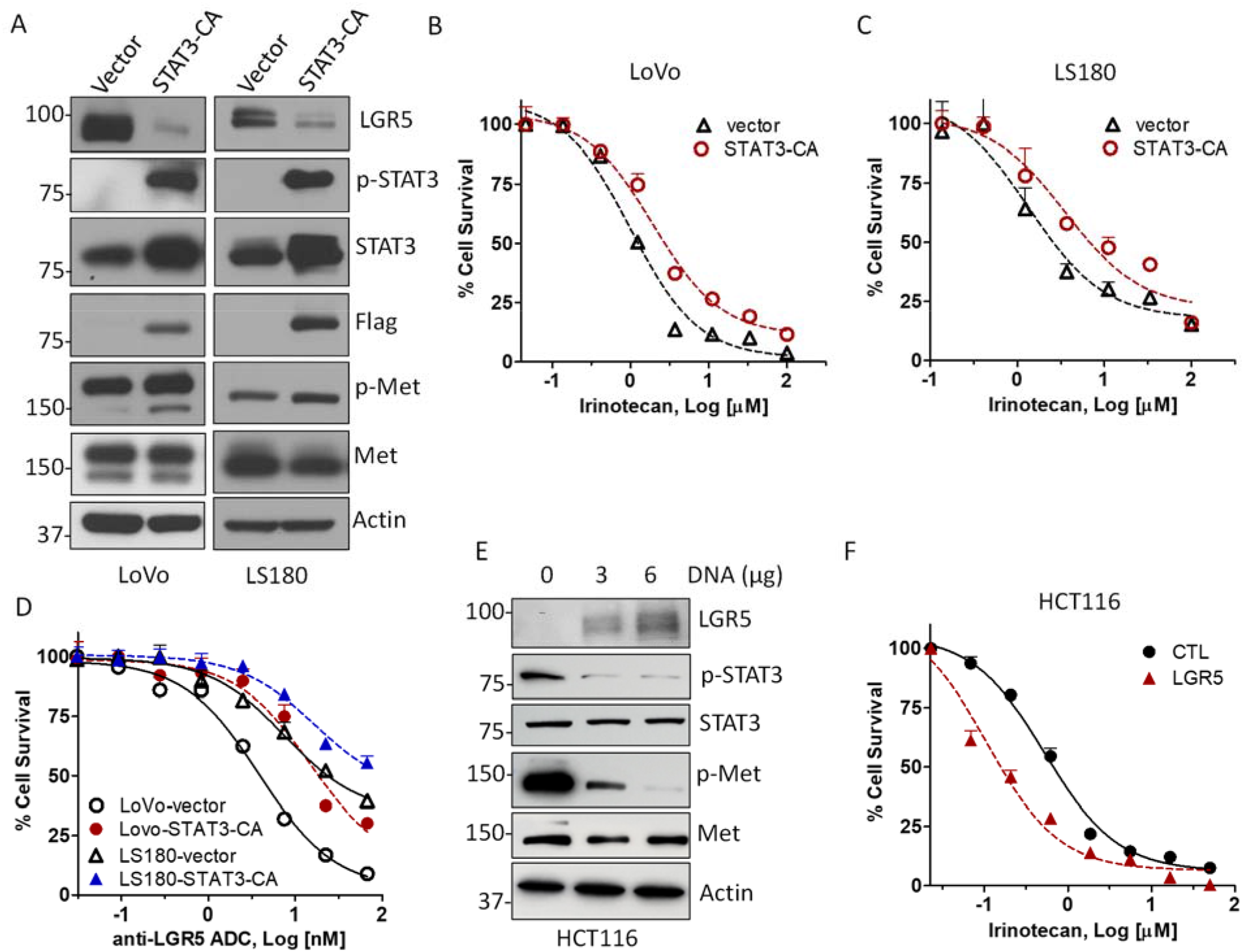
Effects of constitutively active STAT3 and LGR5 overexpression on treatment resistance. (A) Western blot of MET and STAT3 phosphorylation in LoVo and LS180 cells transiently transfected with flag-tagged constitutively active STAT3 (STAT3-CA). (B-C) Cytotoxicity of irinotecan in (B) LoVo and (C) LS180 cells transfected with vector or STAT3-CA after 4 days. (D) Cytotoxicity of anti-LGR5 ADC on CRC cell lines transfected with vector or STAT3-CA after 4 days. (E) Western blot of HCT116 cells transiently transfected with increasing amounts of LGR5 plasmid DNA and effects on MET and STAT3 phosphorylation. (F) Cytotoxicity of irinotecan on HCT116 cells transfected with vector or LGR5 after 4 days. Western blot experiments were performed 3-4 times and cytotoxicity assays were performed 2-3 times in triplicate. Error bars are SEM.

### Drug-induced LGR5^-^ CRC cells have increased MET-STAT3 activation

As gene ablation of LGR5 resulted in increased MET-STAT3 activation, we next evaluated the status of MET-STAT3 in drug-induced LGR5^-^ CRC cells. Treatment with irinotecan or anti-LGR5-ADCs converted LGR5^+^ CRC cells to an LGR5^-^ state over time (Figs. 1A-C, 4A-D). As shown in Figs. 4A-B, increased phosphorylation of MET and STAT3 was detected in LoVo and LS180 cells treated with 10 µM irinotecan. CRC cells treated with 6.5 nM (1 µg/mL) anti-LGR5 ADC also led to increased phosphorylation of MET and STAT3 (Figs. 4C-D). Since LGR5 KD/KO CRC cells have increased levels of active β-catenin, we tested if drug treatment could also modulate β-catenin activation and/or expression. As shown in Figs. 4E-F, neither irinotecan nor ADC treatment significantly altered β-catenin in LoVo cells and a slight reduction in active levels was observed in LS180 cells at the 72-hour time-point. These findings demonstrate that drug-induced loss of LGR5 in CRC cells leads to activation of MET-STAT3, comparable to LGR5 KD/KO CRC cells.

**Figure 4.**
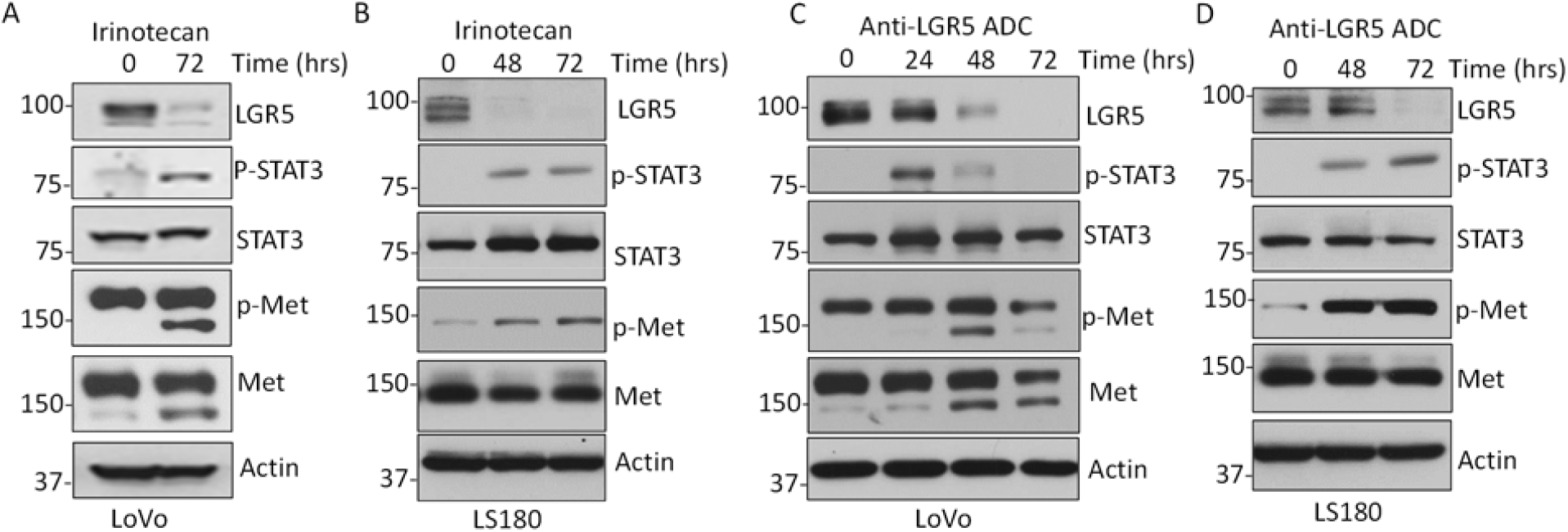
Treatment-induced LGR5-negative CRC cells have increased MET-STAT3 activation. Western blots of MET and STAT3 phosphorylation in (A) LoVo and (B) LS180 cells treated with irinotecan (10µM) at different time-points. (C-D) Western blots of MET and STAT3 phosphorylation in (C) LoVo and (D) LS180 cells treated with anti-LGR5 ADC (6.5 nM) at different time-points. (E-F) Western blots of μ-catenin levels in (E) LoVo and (F) LS180 cells treated with irinotecan (10 μM) or anti-LGR5 ADC (6.5 nM). Experiments were performed at least 3 times.

### LGR5 regulates the sensitivity of CRC cells to MET-STAT3 inhibition

To determine if LGR5 KD/KO CRC cells were more dependent on MET/STAT3 signaling for survival, we tested the effect of MET and STAT3 inhibitors on cell viability. We found that LGR5 KD LoVo and LGR5 KO LS180 cells were more sensitive to crizotinib, stattic, and cryptotanshinone compared to corresponding control cells (Supplementary Figs. S3A-F). As shown in Table 1, the IC_50_ values were approximately 4-to 8-fold lower for LoVo LGR5 KD cells and 1.5-to 6-fold lower for LS180 LGR5 KO cells compared to respective control cells. Since loss of LGR5 also increased levels of active β-catenin (Figs. 2A and 2E), we evaluated the cytotoxic effect of XAV939, a tankyrase inhibitor shown to stimulate β-catenin degradation by stabilizing Axin. However, XAV939 showed no change in efficacy in LGR5 KD/KO cells compared to controls (Supplementary Figs. SG-H). These findings suggest that MET-STAT3 activation may be integral for the enhanced survival capacity of LGR5^-^ CRC cells.

**Table 1.** IC_50_ values for MET and STAT3 inhibitors in CRC cells with and without loss of LGR5 expression. Average IC_50_ values for at least 3 different experiments performed in triplicate. *, P < 0.05; **, P < 0.01; compared to control cell lines by one-way ANOVA.

| CRC cell line | IC<sub>50</sub> (μM) ± SD |  |  |
| --- | --- | --- | --- |
|  | Crizotinib | Cryptotanshinone | Stattic |
| **LoVo** |  |  |  |
| parental | 6.7 ± 1.8 | 6.5 ± 2.9 | 1.2 ± 0.5 |
| shCTL | 6.9 ± 2.2 | 4.2 ± 1.4 | 1.3 ± 0.6 |
| shLGR5-1 | 1.8 ± 0.09* | 1.4 ± 0.7 | 0.2 ± 0.1* |
| shLGR5-2 | 1.5 ± 0.01* | 1.1 ± 0.1 | 0.2 ± 0.1* |
| **LS180** |  |  |  |
| control | 12 ± 5.3 | 11.3 ± 1.8 | 7 ± 1.0 |
| LGR5 KO 1.4 | 7.7 ± 1.5 | 1.9 ± 0.2** | 4.3 ± 0.9* |
| LGR5 KO 1.5 | 5.9 ± 1.7 | 2.4 ± 0.4** | 4.1 ± 0.8* |

### MET-STAT3 inhibitors synergize with irinotecan and ADCs to enhance treatment efficacy in vitro

Given that LGR5 KD/KO CRC cells are more sensitive to MET-STAT3 inhibitors, we next evaluated the efficacy of combination treatment as a strategy to overcome resistance to irinotecan or ADCs. To quantify synergistic effects of the different combinations, combination index (CI) values were calculated using the Chou-Talalay method (17), where CI< 1, =1, or >1 refers to synergistic, additive, or antagonistic activity, respectively (Supplementary Table S1). Synergism was typically observed with increased cytotoxicity, while additive effects or antagonism was observed with lower cytotoxicity. First, we evaluated MET or STAT3 inhibitors, at concentrations below the IC_25_ for control cells, in combination with irinotecan. In LoVo cells, we found that both crizotinib, and stattic synergized with irinotecan (Fig. 6A and Supplementary Table S1). Similar results were observed with combination treatment of cryptotanshinone and irinotecan (Supplementary Fig. S4A). The synergism was much greater in LoVo LGR5 KD cells, as they have increased levels of active MET and STAT3 (Fig. 6B and Supplementary Table S1). Combination treatment of MET or STAT3 inhibitors also synergized with irinotecan in LS180 control and LGR5 KO cells (Figs. 5C-D and Supplementary Table S1), However, the response was less robust in LS180 LGR5 KO cells compared to LoVo LGR5 KD cells (Figs. 5B and 6D). We then tested if MET and STAT3 inhibitors would enhance therapeutic efficacy of anti-LGR5-ADCs. As shown in Figs. 5E-F and Supplementary Table S1, both inhibitors synergized and enhanced efficacy of ADC treatment in LoVo and LS180 cells. However, the effect was less in LoVo cells, which are already highly sensitive to anti-LGR5-MMAE ADCs. Together these data suggest that combination therapy with either MET or STAT3 inhibitors enhances efficacy of chemotherapy and ADCs in CRC cells, likely due to increased activation of MET/STAT3 signaling.

**Figure 5.**
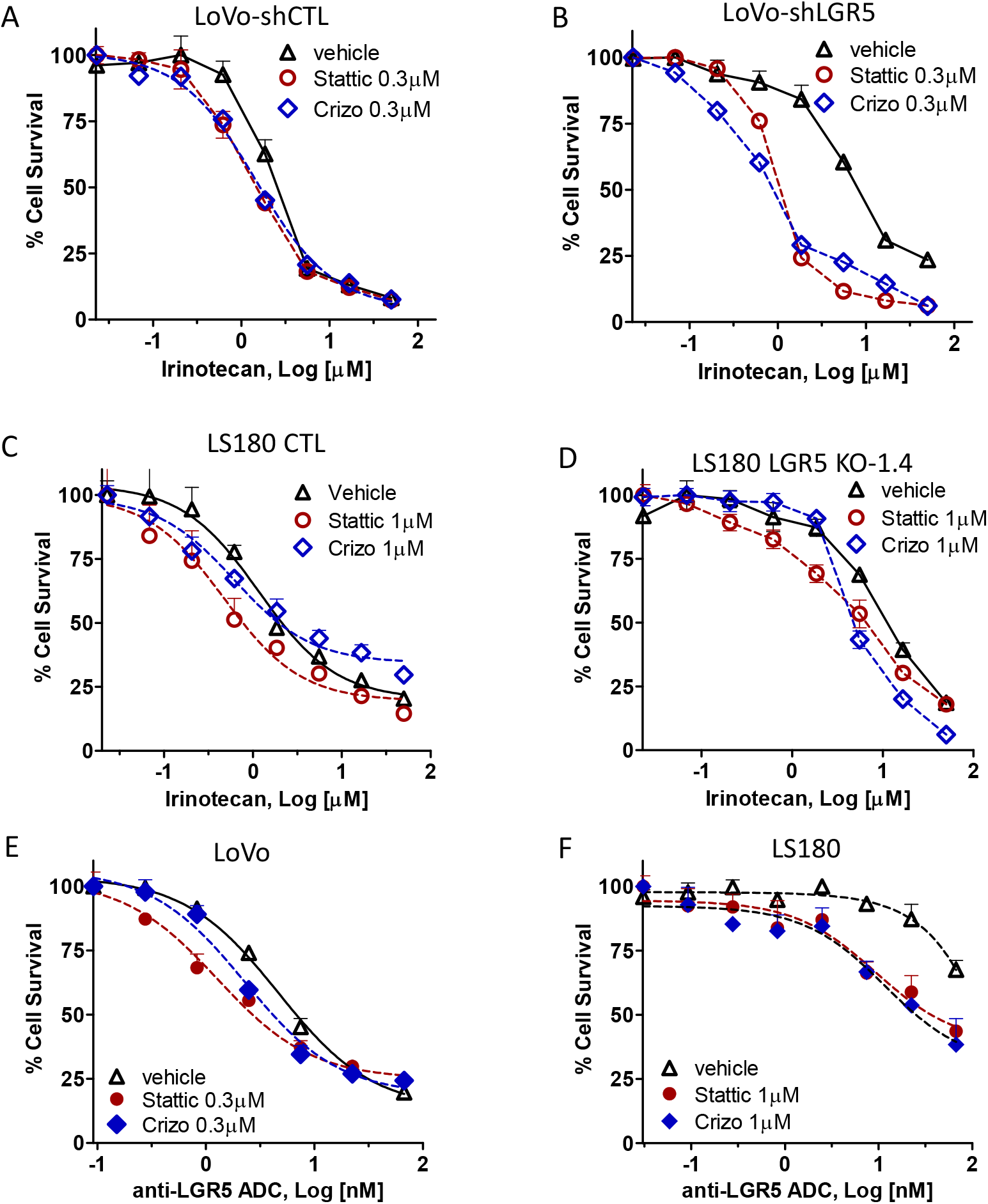
MET-STAT3 inhibitors synergize with irinotecan or ADCs in vitro. (A-B) Cytotoxicity of irinotecan alone or in combination with 0.3 µM crizotinib or stattic in LoVo (A) control and (B) LGR5 KD cells. (C-D) Cytotoxicity of irinotecan alone or in combination with 1 µM crizotinib or stattic in LS180 (C) control and (LGR5 KO cells (E-F) Cytotoxicity of anti-LGR5-MMAE ADCs alone or in combination with crizotinib or stattic in (E) LoVo and (F) LS180 cells. Experiments were performed 2-3 times in triplicates.

**Figure 6.**
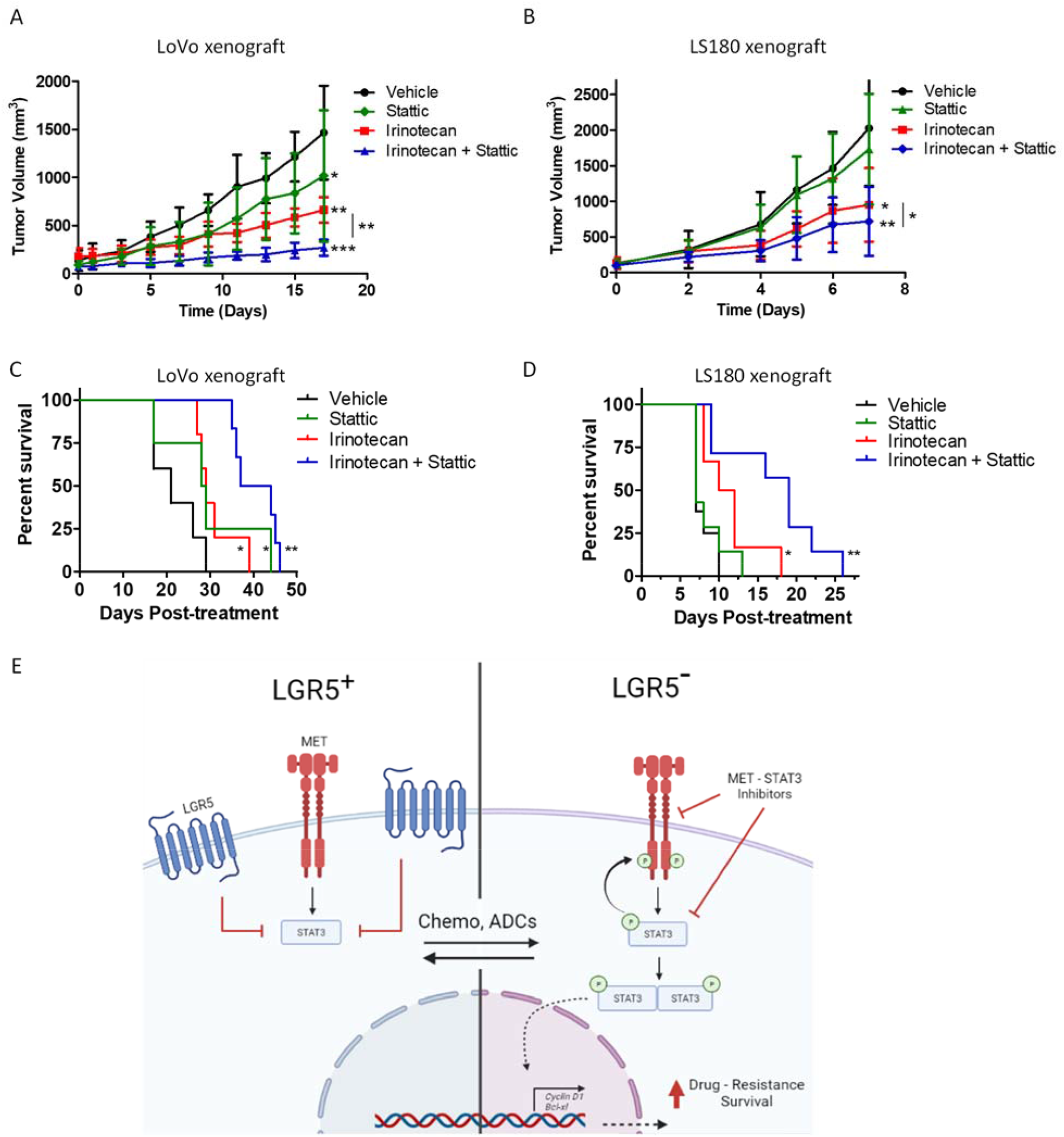
Combination of STAT3 inhibition with irinotecan enhances suppression of tumor growth and increases survival in CRC xenograft models. (A) Tumor growth curve of LoVo xenograft mice treated with vehicle (n=5), 10 mg/kg Stattic (n=4), 20mg/kg irinotecan (n=5), or combination (n=6). (B) Tumor growth curve of LS180 xenograft mice treated with vehicle (n=8), 10 mg/kg Stattic (n=6), 20 mg/kg irinotecan (n=7), or combination (n=7). Statistical analysis was performed using one-way ANOVA and Dunnett’s multiple comparison test. *, P ≤ 0.05, **, P ≤ 0.01, ***, P ≤ 0.001 compared to vehicle unless otherwise indicated. (C-D) Kaplan−Meier survival plot for (C) LoVo and (D) LS180 xenografts. Error bars are SEM and SD for in vitro and in vivo experiments, respectively. (E) Schematic overview of CRC plasticity and MET-STAT3 signaling.

### Combination treatment of irinotecan with a STAT3 inhibitor enhances tumor growth suppression and survival

Since we observed a robust synergistic effect with STAT3 inhibitors in combination with irinotecan, we next evaluated the therapeutic efficacy of this combination using xenograft models. LoVo and LS180 xenograft mice were each randomized into four groups once tumors reached an approximate size of 100-150 mm^3^. Mice were then administered vehicle, 20 mg/kg irinotecan, 10 mg/kg stattic, or irinotecan and stattic in combination (Figs. 6A-B). Although each monotherapy slowed tumor growth in both xenograft models, combination therapy resulted in a more pronounced effect (Figs. 6A-B). Stattic treatment alone led to a 30% and 15% reduction tumor growth in LoVo and LS180 xenografts, respectively. Irinotecan treatment yielded an approximate 53-55% inhibition of tumor growth (Figs. 6A-B). Moreover, combination treatment significantly improved efficacy in both CRC models. Tumor growth inhibition was 82% for LoVo and 65% for LS180 xenografts (Figs. 6A-B). Treatment was terminated after the first animal from each vehicle control group reached the tumor burden limit; however, we continued to monitor survival. As shown in Figs. 6C-D, combination treatment significantly prolonged survival compared to vehicle or monotherapy in both models. No significant effect in body weight nor overt toxicity was observed during treatment (Supplementary Fig. S4B-C). These findings indicate that STAT3-targeted therapy may be highly effective when used in combination with standard chemotherapy to overcome resistance due to plasticity.

## Discussion

In this study, we observed treatment of CRC cells with irinotecan or LGR5-targted ADC led to loss of LGR5 and increased STAT3 activation (Figs. 1 and 5). LGR5 KD/KO CRC cells, shown to be more resistant to standard chemotherapies (15), had increased expression of active STAT3 and STAT3-associated target genes (Figs. 1, 2, and 6). This suggests a potential mechanism whereby LGR5^-^ CRC cells acquire a survival advantage via the STAT3 pathway to promote a more drug resistant phenotype (Fig. 6E). Aberrant activation of STAT3 has been shown in a variety of cancer types including colorectal, prostate, head and neck, breast, liver, and lung cancers (26,27). In CRC, active STAT3 is associated with adverse clinical outcomes (28,29). STAT3 promotes stem cell-like characteristics and regulates diverse cellular processes including survival, proliferation, immune evasion, metastatic potential, and drug resistance (26). As mutation of STAT3 in solid tumors is infrequent, hyperactivation of STAT3 in various cancers typically occurs downstream of various growth factor and cytokine receptors (30).

MET is an RTK that binds to hepatocyte growth factor (HGF) and is highly expressed in CRC. MET has been shown to increase at advanced stages of disease, and contribute to poor prognosis (31,32). Our studies reveal that increased activation of STAT3 in LGR5-negative CRC cells likely occurs downstream of MET (Fig. 2). LGR5 KD/KO cells were more sensitive to MET-STAT3 inhibition compared to control cells, suggesting they are more dependent on STAT3 signaling for survival (Supplementary Fig. S3 and Table 1). Consistently, LGR5 overexpression in CRC cells devoid of endogenous LGR5 led to decreased levels of MET and STAT3 phosphorylation (Fig. 3). As STAT3 inhibitors decreased active MET levels in LGR5 KD CRC cells and STAT3-CA increased MET phosphorylation (Figs. 2-3), this highlights a potential MET-STAT3 feedback mechanism that occurs in CRC. In skin cancer cells, activation of STAT3 led to a slight upregulation in LGR5 expression (33), suggesting interplay between LGR5 and STAT3 may be context dependent. The LGR5-related receptor, LGR4, was shown to be a transcriptional target of STAT3 in osteosarcoma cells and STAT3 KD reduced LGR4 expression (34). LGR4 and LGR5 have different affinities for RSPO1-4 ligands and interact with different Wnt/β-catenin co-receptors and intracellular proteins to regulate distinct functions in cancer cell adhesion/migration, adult stem cells, and during development (10,35-37). Therefore, it is likely that LGR4 and LGR5 may play diverse roles in the regulation of STAT3 and vice versa. STAT3-β-catenin interactions and crosstalk between signaling pathways in CRC and therapy resistance has been reported (38,39). As LGR5 KD/KO increased levels of active β-catenin (Figs. 2A and 2E), this suggests STAT3 may cooperate with β-catenin to promote survival of LGR5^-^ CRC cells. Future studies will be required to elucidate the exact mechanism of how loss of LGR5 leads to MET-STAT3 activation and how STAT3 promotes survival and protects CRC cells from drug-induced cell death. Furthermore, whether MET-STAT3 plays a role in the dynamic plasticity of LGR5+ CSCs during tumor progression and metastasis remains to be elucidated.

In oncogene-addicted cancer cells, including *KRAS* mutant CRC cells, resistance to MEK inhibitors has also been shown to be mediated by STAT3 feedback activation (40-43). One group found MEK inhibition induced MET-STAT3 activity in CRC cells due to inhibition of ADAM17, resulting in decreased shedding of decoy MET (40). Reportedly, MET can undergo ectodomain shedding through the actions of A Disintegrin and Metalloproteinase Domain (ADAM) family members, ADAM10 and ADAM17, and γ-secretases (44,45). The ectodomain has the potential to function as a decoy receptor to block HGF activity and suppress downstream signaling. We observed that both drug treatments and LGR5 KD in LoVo cells led to an increase in what is presumed to be the mature form of MET (145 kDa), suggesting that loss of LGR5 may also lead to activation of a protease(s) that mediates post-translational processing of MET (Figs. 2D, 4A and 4C). LS180 LGR5 KO cells also showed a subtle increase in the ratio of mature MET to pro-MET expression (170 kDa) compared to control cells (Fig. 2E). Numerous other proteases have been implicated in the proteolytic cleavage of MET, resulting in intracellular fragments shown to have either pro-apoptotic or pro-invasive functions (46). The potential role of LGR5 in regulating the proteolytic cleavage and post-translational processing of MET and its relevance in CRC requires further investigation.

Drug combination studies of MET-STAT3 pathway inhibitors with irinotecan or ADCs showed synergism in vitro (Fig. 5). In CRC xenografts, STAT3-targeted therapy in combination with irinotecan significantly enhanced treatment efficacy and overall survival (Fig. 6). Other groups have demonstrated the effectiveness of STAT3 inhibitors on suppression of colorectal tumor growth and sensitization to other chemo- and radiotherapies (47-49). Although STAT3 inhibitors have yet to be approved, several are at various stages of clinical trials for the potential treatment of CRC (26). Our results suggest that combination therapies of MET-STAT3 inhibitors with standard chemotherapy regimens may be more effective than chemotherapy alone to overcome resistance mediated by LGR5^-^ CRC cell populations. Thus, MET-STAT3 inhibition combined with CSC-targeting agents could have promising therapeutic potential and warrants further evaluation.

In summary, we have identified LGR5 as a novel regulator of MET-STAT3 signaling, which may play an intricate role in mediating CRC plasticity. LGR5^+^ CSCs with tumor-initiating capacity have been shown to transition to an LGR5^-^ state to drive metastasis and evade treatment. Therapeutic strategies to target LGR5^+^ CSCs alone have thus far been insufficient to completely eliminate CRC due to tumor heterogeneity and plasticity. This implies that co-targeting both LGR5^+^ and LGR5^-^ CRC cell types may be a more effective treatment for CRC. We reveal that MET-STAT3 activation is triggered when CRC cells transition from an LGR5^+^ to LGR5^-^ state in response to drug treatment or gene ablation. These findings indicate that combination of MET-STAT3 pathway inhibition and irinotecan-based chemotherapy or LGR5-targeted ADCs may offer a promising strategy to target CRC plasticity and improve treatment efficacy and survival.

## Authors’ Contributions

Conceptualization and design: K.S.C; Methodology: K.S.C. and T.A.P., A.N.P., J. J., S.S., L.E.F., and Z. L.; Formal Analysis: K.S.C., T.A.P., J. J., A.N.P.; Writing: K.S.C. and T.A.P; Review and Editing: K.S.C, T.A.P, J.J.; Supervision: K.S.C.

## Acknowledgements

This work was supported by funding from the National Institutes of Health (NCI R01 CA226894) to K.S. Carmon, a fellowship of the Gulf Coast Consortia, on the Training Interdisciplinary Pharmacology Scientists program (T32 GM139801) to J. Jacob, and a Cancer Therapeutics Training Program fellowship (RP210043) to L.E. Francisco. We thank Qingyun Liu and laboratory,Eduardo Vilar-Sanchez, Venkata Lokesh Battula, and Ali Azhdarinia for helpful discussions. Schematic illustration created with BioRender.com

