## Supplementary Information for "Loss of LGR5 through plasticity or gene ablation is associated with therapy resistance and enhanced MET-STAT3 signaling in colorectal cancer cells"

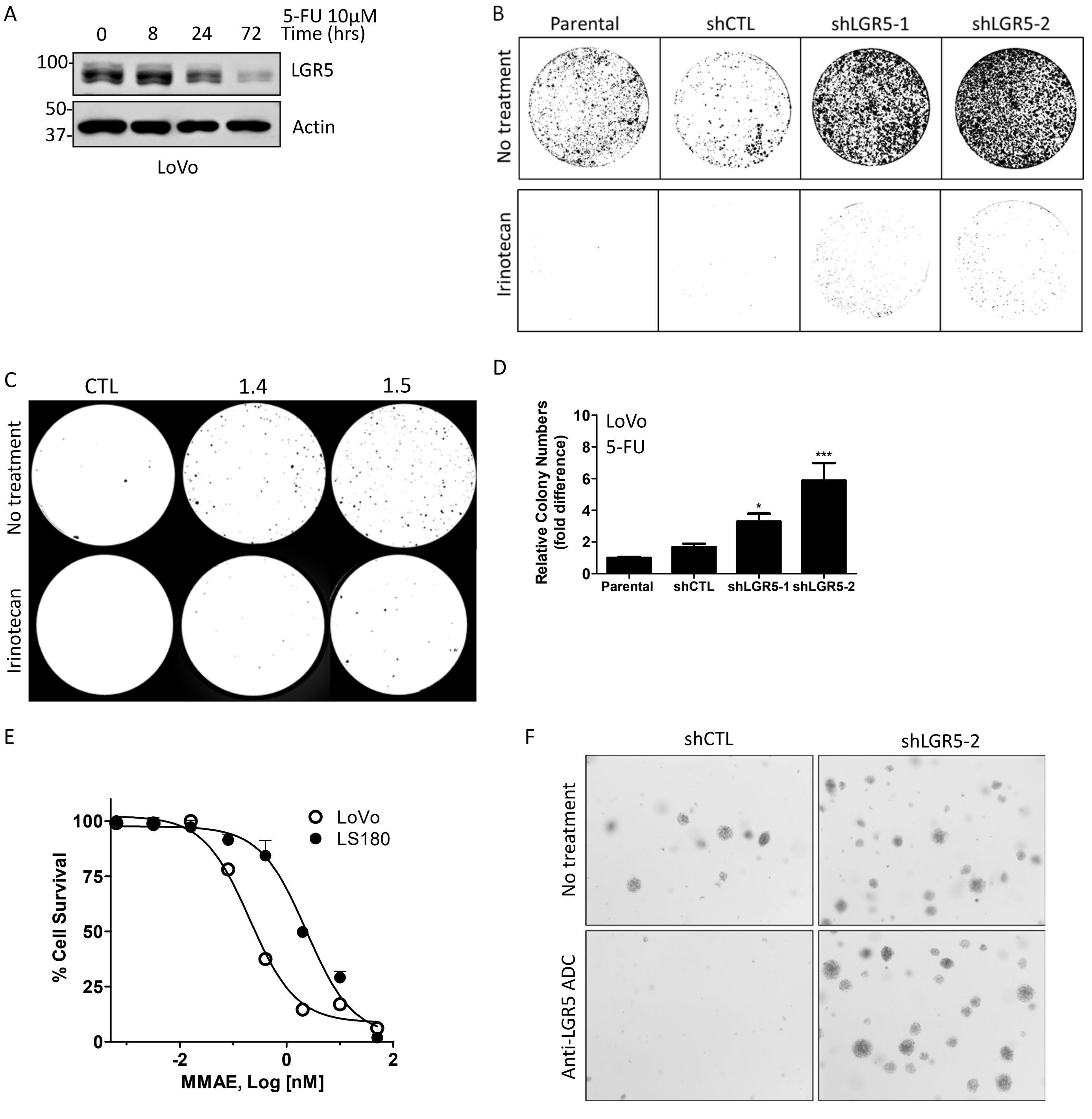


**Supplementary Figure S1. LGR5 expression and clonogenicity.** (A) Western blot analysis of LGR5 expression in LoVo cells treated with 10 µM 5-Fluoruoracil (5-FU). (B-C) Representative images of (B) LoVo and (C) LS180 colony formation assays treated in the presence or absence of 5 µM irinotecan for 1 week (D) Quantitation of LoVo cell colony formation assay in the presence of 10 µM 5-FU. (E) Effect of Irinotecan pretreatment on LoVo cell response to free MMAE. (F) Soft Agar assay of LoVo shCTL and shLGR5 cells in the absence and presence of 6.5 nM anti-LGR5 ADC.

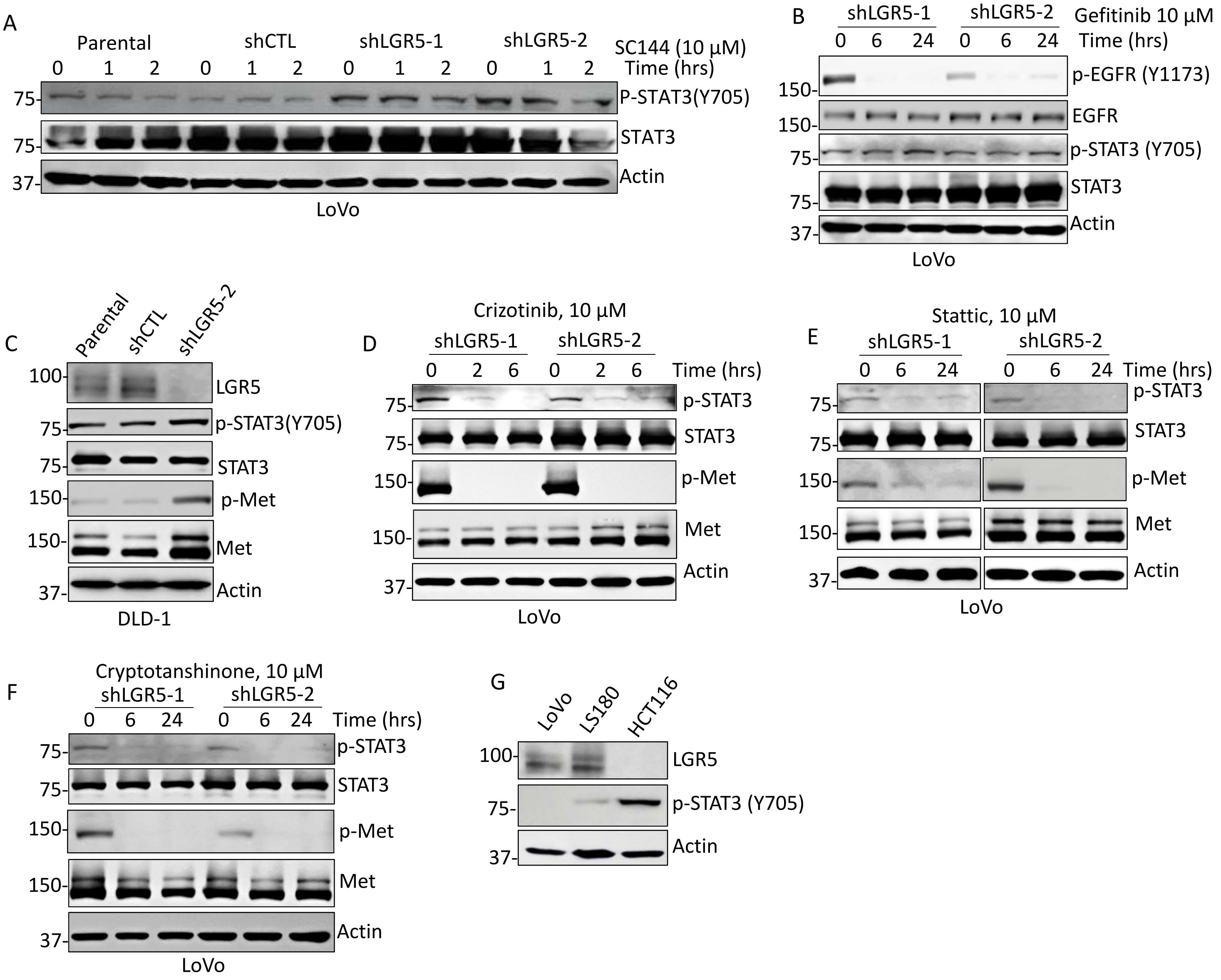


**Supplementary Figure S2. Inhibition of IL6/Gp130, nor EGFR, attenuated STAT3 activation in LGR5 KD/KO cells.** (A) Western blot analysis of STAT3 in LoVo cells treated with 10μM Sc144 for 0, 1, and 2 hours. (B) Western blot analysis of STAT3 and EGFR phosphorylation in LoVo LGR5 KD cells treated with 10μM gefitinib for 0, 6, and 24 hours. (C) Western blot of increased STAT3 and MET phosphorylation in DLD-1 LGR5 KD cells. (D-F) Western blot of inhibitor effects on MET and STAT3 phosphorylation in LoVo LGR5 KD cells treated with (D) crizotinib (10 μM), (E) stattic (10 μM) , or (F) cryptotanshinone (10 μM) at indicated time-points. (G) Western blot of relative LGR5 and p-STAT3 expression in CRC cell lines.

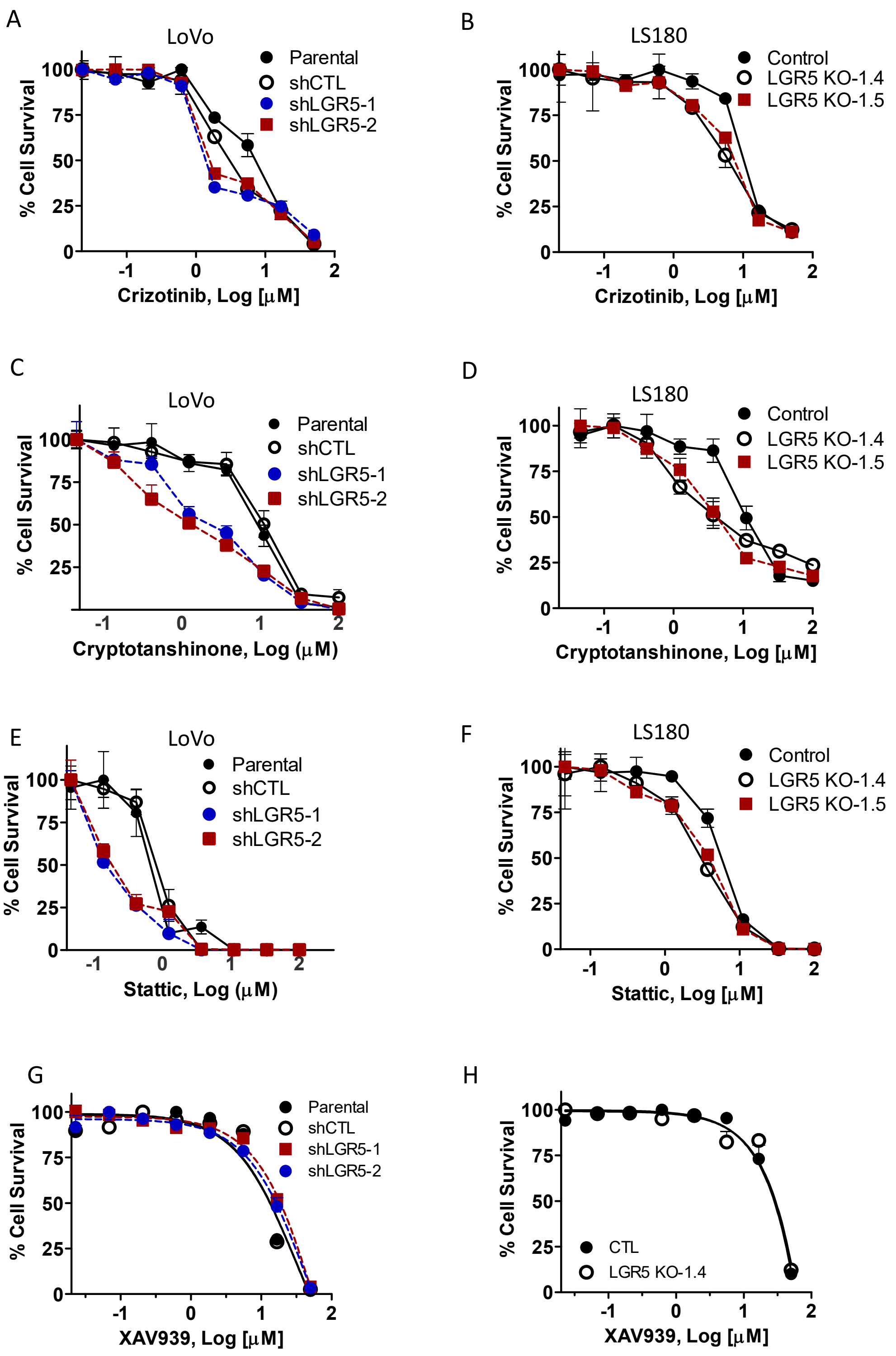


**Supplementary Fig. S3. LGR5 knockdown and knockout cells are more sensitive to inhibitors of MET and STAT3.** (A-B) Cytotoxicity of MET inhibitor, crizotinib in (A) LoVo and (B) LS180 cells. (C-D) Cytotoxicity STAT3 inhibitor, cryptotanshinone in (C) LoVo and (D) LS180 cells. (E-F) Cytotoxicity of STAT3 inhibitor, stattic in (E) LoVo and (F) LS180 cells treated with STAT3 inhibitor, Stattic. Cytotoxicity assays were performed after 4 days using CellTiter-Glo. (G) LoVo and (H) LS180 cells treated with Wnt signaling inhibitor, XAV939. Cytotoxicity assays were performed after 4 days using CellTiter-Glo. Experiments were performed 3 times in triplicate. Error Bars are SEM.

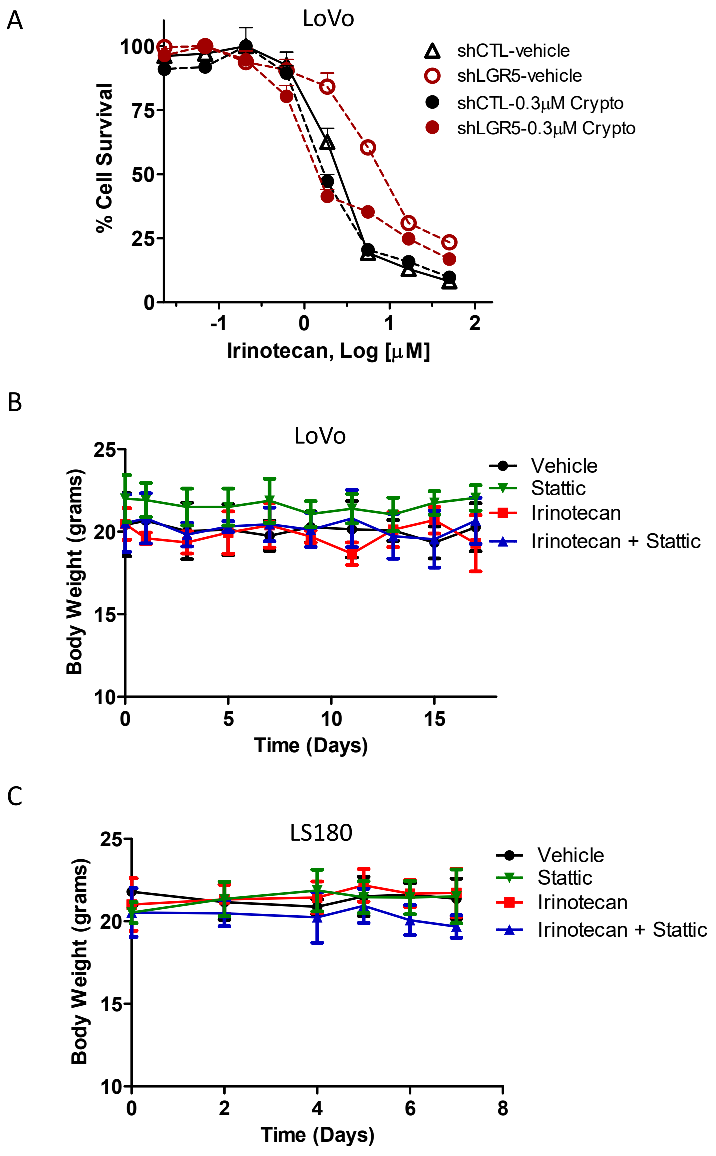


**Supplementary Figure S4. Combination therapy and body weight of CRC xenograft mice.** (A) Cytotoxicity of irinotecan in combination with 0.3 μM cryptotanshinone in LoVo control and LGR5 KD cells. (B-C) Average body weight of (B) LoVo and (C) LS180 xenograft mice over the course of treatment. Error bars are SD.

| Combination Index (CI) values | | | | | | | | |
|---|---|---|---|---|---|---|---|---|
| | LoVo shCTL | | LoVo shLGR5 | | LS180 CTL | | LS180 LGR5 KO | |
| Irinotecan (µM) | Stattic (0.3 µM) | Crizotinib (0.3 µM) | Stattic (0.3 µM) | Crizotinib (0.3 µM) | Stattic (1 µM) | Crizotinib (1 µM) | Stattic (1 µM) | Crizotinib (1 µM) |
| 17 | 0.87 | 1.05 | 0.22 | 0.42 | 0.99 | 1.72 | 0.86 | 0.44 |
| 6.0 | 0.42 | 0.59 | 0.11 | 0.28 | 0.65 | 0.97 | 0.87 | 0.50 |
| 2.0 | 0.39 | 0.60 | 0.09 | 0.18 | 0.51 | 0.76 | 0.77 | 1.63 |
| 0.6 | 0.40 | 0.79 | 0.29 | 0.28 | 0.49 | 0.74 | 0.93 | 1.49 |
| 0.2 | 1.70 | 1.24 | 1.76 | 0.37 | 0.65 | 0.93 | 1.39 | 1.08 |
| ADC (nM) | (LoVo) | | | | (LS180) | | | |
| 65 | 2.25 | 2.57 | ND | ND | 0.18 | 0.11 | ND | ND |
| 20 | 1.18 | 1.19 | ND | ND | 0.22 | 0.13 | ND | ND |
| 7.0 | 0.77 | 0.78 | ND | ND | 0.24 | 0.16 | ND | ND |
| 2.0 | 0.72 | 0.82 | ND | ND | 0.46 | 0.30 | ND | ND |
| 0.7 | 0.68 | 1.00 | ND | ND | 0.40 | 0.26 | ND | ND |

**Supplementary Table S1. Combination index (CI) values for irinotecan or ADCs in combination with MET or STAT3 inhibitors.** Combination index values determined by the Chou-Talalay method using CompuSyn. Synergism defined as CI<1 (Red), additive effect as CI=1 (White), and antagonism as CI>1 (Blue).
